# “*Candidatus* Dechloromonas phosphatis” and “*Candidatus* Dechloromonas phosphovora”, two novel polyphosphate accumulating organisms abundant in wastewater treatment systems

**DOI:** 10.1101/2020.11.05.369777

**Authors:** Francesca Petriglieri, Caitlin Singleton, Miriam Peces, Jette F. Petersen, Marta Nierychlo, Per H. Nielsen

## Abstract

Members of the genus *Dechloromonas* are often abundant in enhanced biological phosphorus removal (EBPR) systems and are recognized putative polyphosphate accumulating organisms (PAOs), but their role in phosphate (P) removal is still unclear. Here, we used 16S rRNA gene sequencing and fluorescence *in situ* hybridization (FISH) to investigate the abundance and distribution of *Dechloromonas* spp. in Danish wastewater treatment plants. Two species were abundant, novel, and uncultured, and could be targeted by existing FISH probes. Raman microspectroscopy of probe-defined organisms (FISH-Raman) revealed the levels and dynamics of important intracellular storage polymers in abundant *Dechloromonas* spp. in the activated sludge from four full-scale EBPR plants and from a lab-scale sequencing batch reactor fed with different carbon sources (acetate, glucose, glycine, and glutamate). Moreover, 7 distinct *Dechloromonas* species were determined from a set of 10 high-quality metagenome-assembled genomes (MAGs) from Danish EBPR plants, each encoding the potential for poly-P, glycogen, and polyhydroxyalkanoates (PHA) accumulation. The two most abundant species exhibited an *in situ* phenotype in complete accordance with the metabolic information retrieved by the MAGs, with dynamic levels of poly-P, glycogen, and PHA during feast-famine anaerobic-aerobic cycling, legitimately placing these microorganisms among the important PAOs. As no isolates are available for the two species, we propose the names *Candidatus* Dechloromonas phosphatis and *Candidatus* Dechloromonas phosphovora.

## Introduction

The enhanced biological phosphorus removal (EBPR) process is becoming globally popular in wastewater treatment. In EBPR, phosphorus (P) removal can be achieved without chemical addition and the P-enriched biomass can be used for P recovery or as agricultural fertilizer [1]. The EBPR process depends on the ability of polyphosphate accumulating organisms (PAOs) to store polyphosphate (poly-P) as an intracellular storage compound when exposed to alternating carbon-rich anaerobic (feast) and carbon-deficient aerobic (famine) conditions. The metabolism of well-described PAOs, such as *Ca*. Accumulibacter and *Tetrasphaera*, has been extensively investigated in lab-scale enrichments [2–6] and recently verified in full-scale EBPR plants too [7]. While *Ca*. Accumulibacter cycles poly-P during dynamic feast-famine conditions with PHA and glycogen storage, *Tetrasphaera* spp. exhibit a different behavior, as no PHA or glycogen have been identified *in situ* [7].

Several putative PAOs have been identified through the detection of intracellular poly-P [8–11], including the genus *Dechloromonas*, first described in EBPR plants by Goel et al. [12]. Over recent years, *Dechloromonas* has been consistently found in high abundances in many EBPR plants in Japan [13], China [14], Denmark [15], and worldwide [1]. *Dechloromonas* spp. were originally isolated as aromatic compound-degrading bacteria and facultative anaerobic nitrate-reducing bacteria [16, 17]. Since then, several *Dechloromonas* populations have been shown to accumulate poly-P and PHA *in situ* and/or in axenic cultures [11, 13], and several studies have also suggested their potential role as PAOs [18–20]. Furthermore, analysis of the *Dechloromonas aromatica* strain RCB genome revealed the genes required for poly-P accumulation, including polyphosphate kinase, and exopolyphosphatase [21]. *Dechloromonas* spp. likely also have important nitrogen cycling roles in full-scale EBPR plants, as uncultured *Dechloromonas* has the ability to respire with NO3- and/or NO2-*in situ* [11, 22, 23], and are suggested to be denitrifying PAOs [19, 20]. Other uncultured members of the genus *Dechloromonas* have also shown the potential for a glycogen accumulating organism (GAO) phenotype [22, 24], long considered competitors of PAOs. The genus encompasses a broad range of metabolisms integral to the wastewater treatment ecosystem, exposing the need for a deeper characterization of these organisms.

Recent advances in high-throughput sequencing methods have enabled the creation of comprehensive ecosystem-specific full-length 16S rRNA gene databases [25]. These databases can be used for detailed phylogenetic analyses, facilitating the design of specific fluorescence *in situ* hybridization (FISH) probes and re-evaluation of existing ones for the identification and characterization of key species [26]. Moreover, it is now possible to assemble high-quality, near-complete genomes from deeply sequenced metagenomes, revealing the metabolic potential of these organisms [26–28]. However, *in situ* validation of the genomic potential is critical and required to confirm their predicted role in the system. Recently, Raman microspectroscopy combined with FISH was used to identify and quantify intracellular storage compounds in *Ca*. Accumulibacter and *Tetrasphaera* [7]. While these lineages accounted for 24–70% of total P removal [7], undescribed PAOs likely remove a large portion of P. Identifying these novel PAOs and obtaining more insights into their physiology is essential to improve the management and efficacy of resource recovery in EBPR treatment plants.

In this study, we present and characterize novel uncultured species belonging to the genus *Dechloromonas* and show that they are actively involved in P removal in full-scale EBPR plants. We confirmed the presence and dynamics of intracellular poly-P and other storage compounds in the novel species using the new Raman-based approach on activated sludge from full-scale WWTPs. The effect of several individual substrates on P-release was tested using sludge from a lab-scale sequencing batch reactor (SBR). Moreover, the genes for PAO metabolism were identified in 10 high-quality *Dechloromonas* metagenome-assembled genomes (MAGs) from Danish EBPR plants. The combination of genome-based investigations and *in situ* analyses provides the first detailed insight into the ecophysiology of these abundant, widespread, and novel PAOs.

## Materials and Methods

### Full-scale activated sludge batch experiments for P cycling

Batch experiments were conducted on fresh activated sludge to analyze the poly-P-content per cell of FISH-identified *Dechloromonas* cells under anaerobic and aerobic conditions. Fresh samples were collected from four large full-scale Danish WWTPs (Lynetten, Ejby Mølle, Viby, and Aalborg West) and aerated for 30 min to exhaust most intracellular carbon source reserves. After aeration, sludge was transferred to serum bottles and sealed with a butyl septum and aluminium cap. A substrate solution containing acetate, glucose, and casamino acids was added, with a final concentration of the three components of 500, 250, and 250 mg L^-1^, respectively. Ultrapure nitrogen was used to flush the headspace in each bottle to ensure anaerobic conditions. The serum bottles were kept at room temperature (∼22°C) with shaking for 3 h. Samples for ortho-P analysis were taken every 20 min for the first hour of the experiment, and every 30 min during the remaining 2 h. Initial samples (0 h) and at the end of the experiment (3 h) were fixed for FISH and Raman analyses (see below).

### P cycling experiments in SBR reactor

A sequencing batch reactor (SBR) with 5 L working volume was operated with 8 h cycles based on Marques et al., [2]. Each cycle included a 4 h anaerobic phase, a 1 h settling/decant phase, and a 3 h aerobic phase. The SBR was operated with a hydraulic retention time of 13 h, a solid retention time of 15 d, organic loading rate of 0.6 g COD L^-1^ d^-1^ and phosphate loading rate of 0.2 g P L^-1^ d^-1^. To maintain anaerobic conditions, nitrogen was bubbled continuously at a flow rate of 4 L min^-1^. The aerobic phase was controlled at oxygen saturation point (∼9 mg L^-1^) by bubbling compressed air at a flowrate of 4 L min^-1^. Temperature was controlled at 19 ± 1°C. pH was controlled at 7.5 by dosing HCl (0.5 M) or NaOH (0.4 M). During aerobic and anaerobic phase, the SBR was completely stirred using an overhead stirrer set at 400 rpm. The SBR was fed with a synthetic medium containing casein hydrolysate (4.4 g L^-1^; 5.5 g COD L^-1^) (Sigma-Andrich, USA) during the anaerobic phase in 6 pulses every 30 min (30 mL/pulse). After the decanting phase, during the first 5 min of the aerobic phase, the SBR was fed (3.1 L) with mineral medium containing per litre: 317 mg K_2_HPO4, 190 mg KH_2_PO_4_, 123 mg NH_4_Cl, 198 mg MgSO_4_·7H_2_O, 92 mg CaCl_2_·2H_2_O, 1.7 mg N-Allylthiourea (ATU), 6.7 mg ethylenediaminetetraacetic (EDTA), and 0.7 mL of micronutrient solution. The micronutrient solution was based on Smolders et al., [29] and contained per litre: 1.5 g FeCl_3_·6H_2_O, 0.15 g H_3_BO_3_, 0.03 g CuSO_4_·5H_2_O, 0.18 g KI, 0.12 g MnCl_2_·_4_H_2_O, 0.06 g Na_2_MoO_4_·2H_2_O, 0.12 g ZnSO_4_·7H_2_O, and 0.15 g CoCl_2_·6H_2_O.

Batch tests were performed to evaluate the phosphate release from different carbon sources. Biomass from the SBR was harvested after the aerobic phase (i.e., after phosphate uptake) and directly used for the batch test without amendments. 25 mL of biomass was dispensed in 30 mL serum bottles and sealed with butyl septa and aluminium caps. The serum bottles were flushed with nitrogen (4 L min^-1^) for 15 min to ensure anaerobic conditions. Five carbon sources (casein hydrolysate, acetate, glutamate, glycine, or glucose) were tested individually. The cycle started with the addition of 200 μL of stock solution to an initial cycle concentration of 200 mg COD L^-1^. The anaerobic cycle lasted for 4 h. The bottles were mixed by manual swirling before each sampling event. Blank controls without carbon source addition were used to evaluate the endogenic phosphate release. All the experiments were performed in duplicate. Samples from the start (0 h) and the end of the experiment (4 h) were fixed for FISH and Raman analyses.

### Chemical analyses

The ortho-P release into the liquid phase was analysed in accordance with ISO 6878:2004 using the ammonium molybdate-based colorimetric method. Samples for total P measurement were taken at the start of the P-release experiment. Sludge collected from the bottles was homogenized and stored at −20°C until further analysis. 67% nitric acid was used to dissolve 0.5 mL of each sludge sample and the samples were microwave heated according to U.S. EPA (2007). The total amount of P in the samples was analysed by inductively coupled plasma optical emission spectrometry (ICP-OES) in accordance with Jørgensen et al., [30].

### Sampling and fixation

Biomass samples from SBR batch reactors and full-scale activated sludge were either stored at −80°C for sequencing workflows or fixed for FISH with 96% ethanol or 4% PFA, as previously described [31] and stored at −20°C until analysis.

### DNA extraction

DNA extraction of activated sludge samples from the MiDAS collection [32] was performed as described by Stokholm-Bjerregaard et al. [15]. Briefly, DNA was extracted using the FastDNA spin kit for soil (MP Biomedicals), following the manufacturer’s indications, but with an increase of the bead beating to 6 m/s for 4×40 s, using a FastPrep FP120 (MP Biomedicals).

### Community profiling using 16S rRNA gene amplicon sequencing

Amplicon sequence variant (ASV) analysis is described in Dueholm et al. [25] and Nierychlo et al [33]. Community profiling of Danish WWTPs was performed using 16S rRNA gene amplicon sequencing data collected from 2006 to 2018 from 27 different Danish WWTPs, as part of the MiDAS project [32]. Data was analyzed using R (version 3.5.2) (R Development Core Team, 2018), RStudio software [34] and visualized using ampvis2 [35] and ggplot2 [36].

### Phylogenetic analysis and FISH probes evaluation

Phylogenetic analysis of 16S rRNA gene sequences and evaluation of existing FISH probes for the genus *Dechloromonas* were performed using the ARB software v.6.0.6 [37]. A phylogenetic tree was calculated based on the full-length 16S rRNA gene sequences retrieved from the MiDAS 3.7 database [25] and Silva 138 SSURef Nr99 [38], using the maximum likelihood method and a 1000-replicate bootstrap analysis. Coverage and specificity of the existing probes Bet135 [11] and Dech443 [22] were evaluated and validated *in silico* with the MathFISH software for hybridization efficiencies of target and potentially weak non-target matches [39]. All probes were purchased from Sigma-Aldrich (Denmark) or Biomers (Germany), labelled with 6-carboxyfluorescein (6-Fam), indocarbocyanine (Cy3), or indodicarbocyanine (Cy5) fluorochromes.

### Fluorescence in situ hybridization (FISH) and quantitative FISH (qFISH)

FISH was performed on full-scale activated sludge samples as well as sludge from SBR reactor as described by Daims et al. [40]. Optimal formamide concentration and use of competitors or helper probes was applied as recommended by the authors for each probe [11, 22]. Quantitative FISH (qFISH) biovolume fractions of individual *Dechloromonas* spp. were calculated as a percentage area of the total biovolume, hybridizing the EUBmix probes, that also hybridizes with the specific probe. qFISH analyses were based on 30 fields of view taken at 630× magnification using the Daime image analysis software [41]. Microscopic analysis was performed with a white light laser confocal microscope (Leica TCS SP8 X).

### Raman microspectroscopy

Raman microspectroscopy was applied in combination with FISH as previously described [7]. Briefly, FISH was conducted on optically polished CaF2 Raman slides (Crystran, UK). *Dechloromonas*-specific (Cy3) probes [11, 22] were used to locate the target cells for Raman analysis. After bleaching, 100 spectra from single-cells were obtained using a Horiba LabRam HR 800 Evolution (Jobin Yvon – France) equipped with a Torus MPC 3000 (UK) 532 nm 341 mW solid-state semiconductor laser. The Raman spectrometer was calibrated prior to obtaining all measurements to the first-order Raman signal of Silicon, occurring at 520.7 cm^-1^. The incident laser power density on the sample was attenuated down to 2.1 mW/μm^2^ using a set of neutral density (ND) filters. The Raman system is equipped with an in-built Olympus (model BX-41) fluorescence microscope. A 50X, 0.75 numerical aperture dry objective (Olympus M Plan Achromat-Japan), with a working distance of 0.38 mm, was used throughout the work. A diffraction grating of 600 mm/groove was used and the Raman spectra collected spanned the wavenumber region of 200 cm^-1^ to 1800 cm^-1^. The slit width of the Raman spectrometer and the confocal pinhole diameter were set respectively to 100 μm and 72 μm. Raman spectrometer operation and subsequent processing of spectra were conducted using LabSpec version 6.4 software (Horiba Scientific, France). All spectra were baseline corrected using a 6^th^ order polynomial fit.

### Identification, annotation, and metabolic reconstruction of Dechloromonas species

A set of 1083 high-quality MAGs were searched for *Dechloromonas* spp. based on taxonomic classification using GTDB-Tk [42] v1.0.2 [26]. Distinct species were determined using dRep [43] v2.3.2 and 95% ANI genome clustering. The MAGs were annotated with KEGG orthology [44] numbers using EnrichM (https://github.com/geronimp/enrichM) v0.5.0 against the uniref100 database, with pathways identified based on genes and modules defined by KEGG. Pathways were considered complete if 100% of the genes in the KEGG module were identified. For *Ca*. D. phosphovora, the species was presented as encoding the pathway in Figure 6 if at least one of the three MAGs encoded the full pathway, see STable1 and SDataFile1 for details. The 16S, 23S, 5S rRNA, and tRNA genes were identified using Prokka [45] v1.14 and Infernal [46] v1.1.2. Fxtract (https://github.com/ctSkennerton/fxtract) v2.3 extracted the 16S rRNA gene sequences for placement in the phylogenetic 16S rRNA gene tree. The phylogenetic genome tree of *Dechloromonas* was created using GTDB-Tk v1.0.2 and the concatenated alignment of 120 single copy proteins. The concatenated protein alignments created by GTDB-Tk were used as input into IQ-TREE v2.0 [47] using the WAG+G model and 100 bootstrap iterations, with 5 *Ca*. Accumulibacter genomes used as an outgroup.

## Results and discussion

### Diversity and distribution of *Dechloromonas* spp. in full-scale wastewater treatment plants

Phylogenetic analysis based on full-length high quality 16S rRNA gene sequences retrieved from 24 Danish WWTPs [25] revealed the presence of 12 *Dechloromonas* species (Figure 1). Some of them were well-known from axenic cultures, such as *Dechloromonas denitrificans* [17], *Dechloromonas hortensis* [48], and *Dechloromonas agitata* [16], while the majority were novel and undescribed.

**Figure 1.**
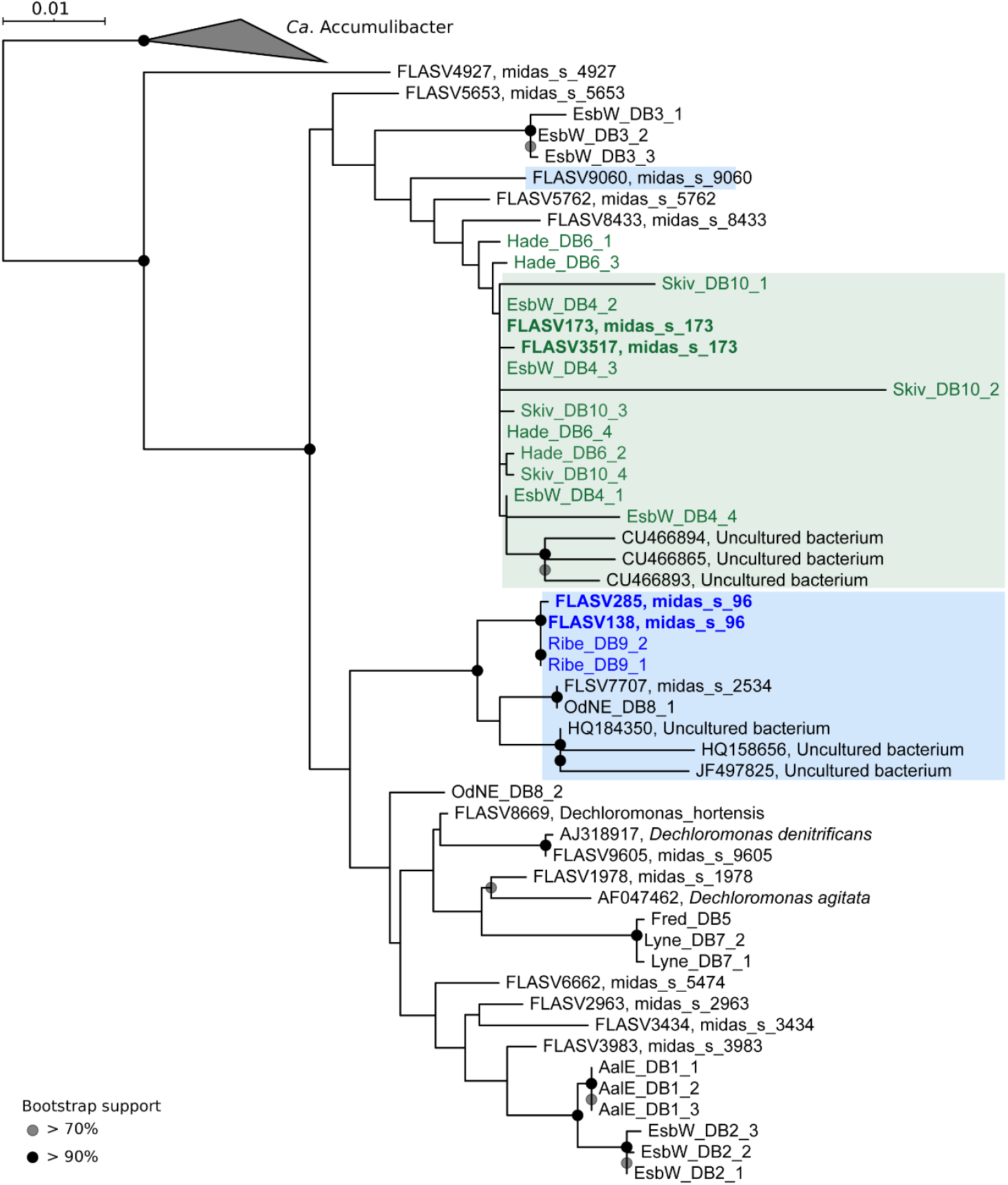
Maximum-likelihood (PhyML) 16S rRNA gene phylogenetic tree of the genus *Dechloromonas* including full-length sequences from MiDAS3 database (FLASVxxx), MAG (xxxx_DBx_x) and Silva138 SSURef Nr99 (indicated by sequence accession numbers). Colours indicate the two most abundant *Dechloromonas* species in activated sludge (*Ca*. D. phosphovora in green, midas_s_173, and *Ca*. D. phosphatis in blue, midas_s_96, shown in bold) and their related MAG 16S rRNA gene sequences. Coverage of FISH probes Dech443 and BET135 is shown as green- and blue-shaded area, respectively. Bootstrap values from 1000 re-samplings are indicated for branches with >70% (grey circle), and >90% (black) support. The scale bar represents substitutions per nucleotide base.

A survey of 20 Danish WWTPs using data from the MiDAS project [33] showed that *Dechloromonas* was the second most abundant genus among well-recognized and putative PAOs (Figure 2A). This high abundance is in accordance with our recent study of PAOs and putative PAOs across the world in 101 EBPR plants [1]. The genus constituted on average 2.6% of the total reads across all Danish plants with abundances reaching 20% in some samples. The two most abundant species, midas_s_96 and midas_s_173 (Figure 2B), for which we propose the names *Candidatus* Dechloromonas phosphatis and *Candidatus* Dechloromonas phosphovora, were targeted by the existing FISH probes Bet135 [11] and Dech443 [22], respectively, with good specificity and coverage (Figure 1). ASV85 was classified to *Ca*. D. phosphovora with the aid of 16S rRNA genes retrieved from the MAGs (see below), indicating its potentially higher abundance in some WWTPs. Application of the probes to fixed activated sludge biomass showed rod-shaped cells (1.4×1.2 µm and 1.2×0.8 µm, respectively), often arranged in spherical microcolonies (Figure 3).

**Figure 2.**
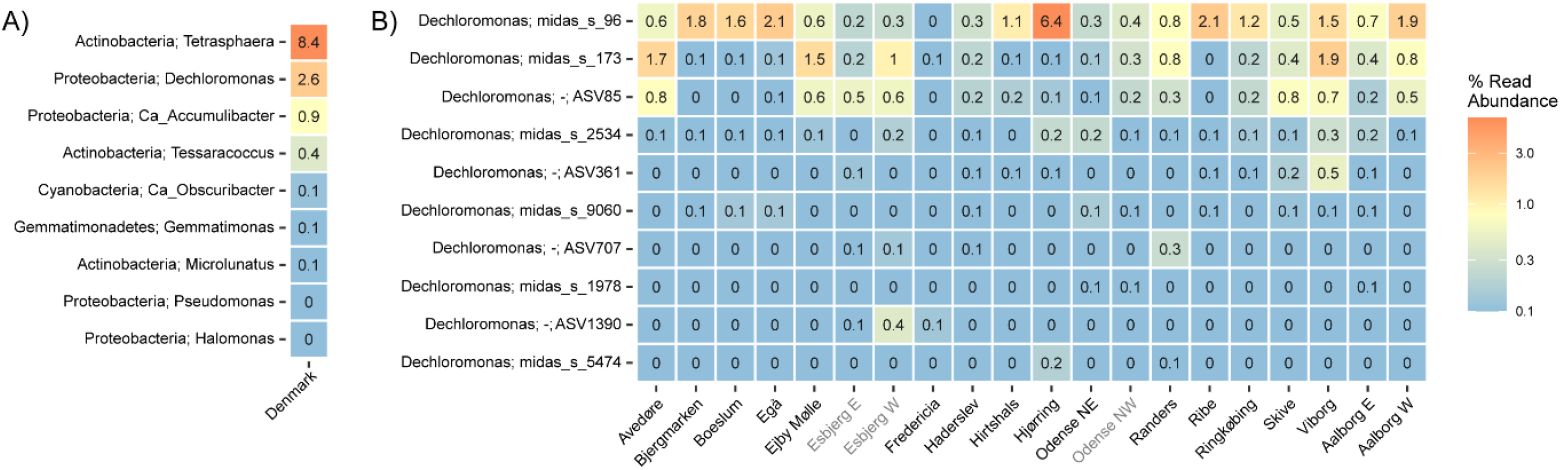
A) Abundance of known or putative PAOs in Denmark (phylum and genus name is given). B) Abundance of *Dechloromonas* species in Danish WWTPs. ASV numbers represent taxa that could not be confidently classified at species level, as indicated by “-”. midas_s_96 corresponds to *Ca*. D. phosphatis and midas_s_173 and ASV85 to *Ca*. D. phosphovora. Names of non-EBPR plants are shown in grey.

**Figure 3.**
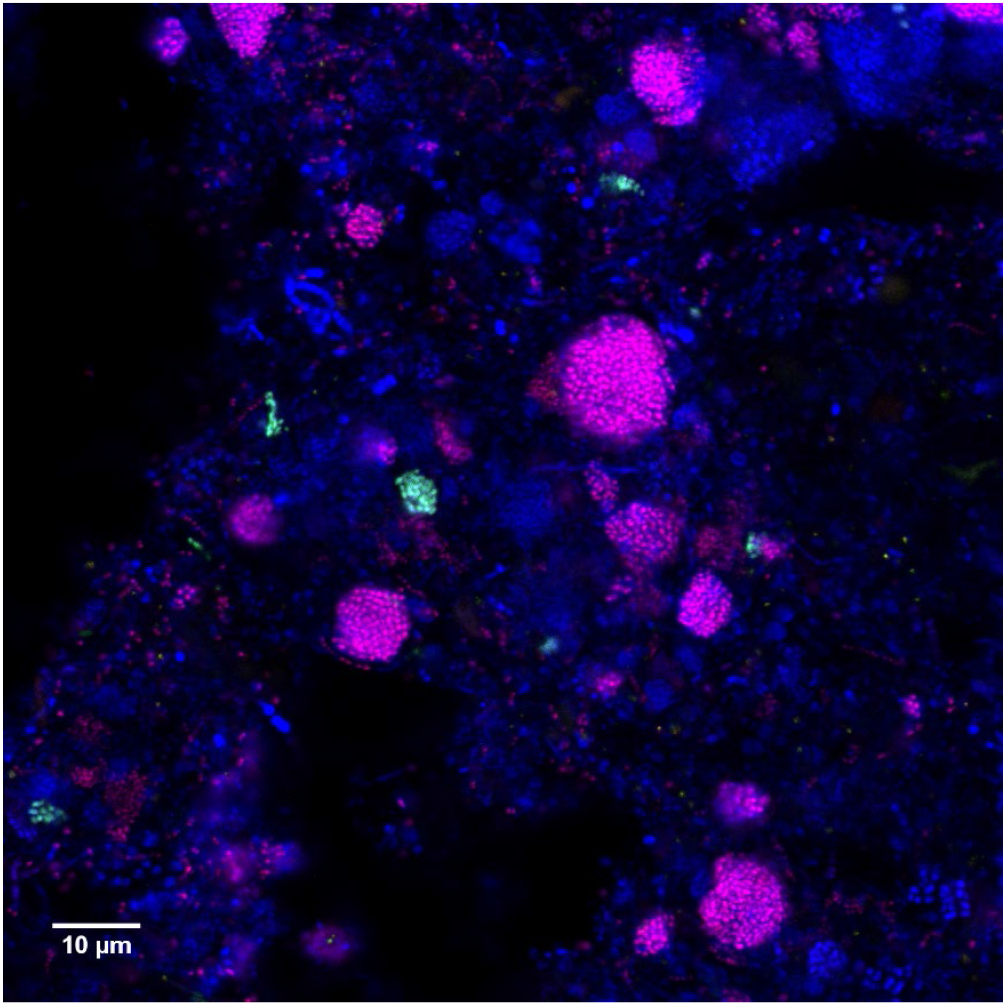
Composite FISH micrographs of the two most abundant *Dechloromonas* species in Danish WWTPs. Specific probes Bet135 (Cy3-label, red) and Dech443 (FLUOS, green) are targeting the *Ca*. D. phosphatis and *Ca*. D. phosphovora, respectively. All bacteria are targeted by EUBmix probe (Cy5-label, blue) and overlap appears in magenta (*Ca*. D. phosphatis - Bet135) or cyan (*Ca*. D. phosphovora - Dech443).

Quantitative FISH analyses confirmed the relative abundances obtained by 16S rRNA gene amplicon sequencing of selected samples (Table 1), showing little differences, most likely due to the presence of several 16S rRNA gene copies in some of the species [21].

**Table 1.**
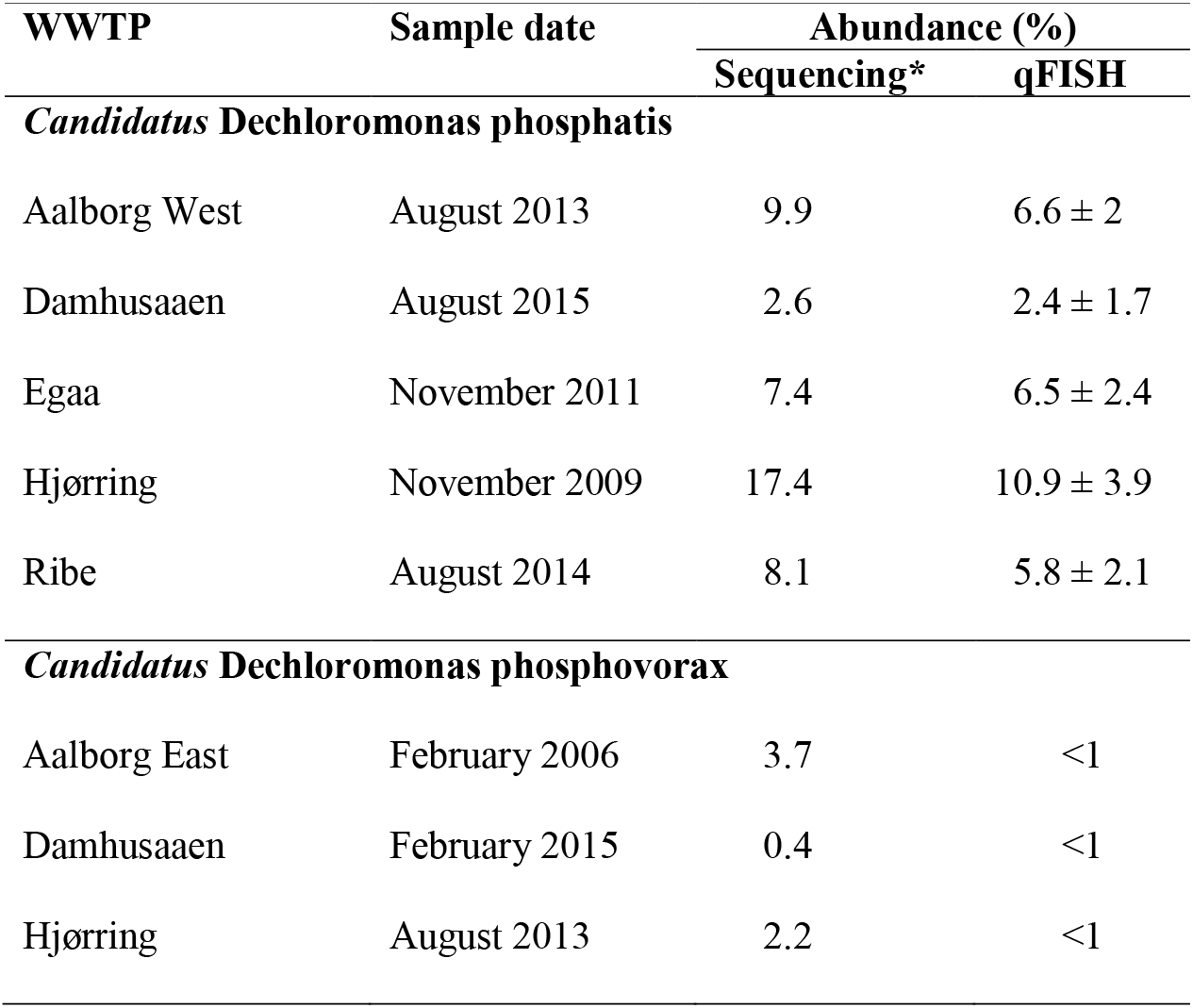
Abundance estimation: 16S rRNA amplicon sequencing and qFISH (percentage of total) of full-scale activated sludge samples.

### *In situ* quantification of storage polymers in *Dechloromonas* spp. in full-scale EBPR plants

In order to identify, quantify, and explore the dynamics of intracellular storage polymers, such as poly-P, PHA, and glycogen, in the *Dechloromonas* spp., we performed anaerobic-aerobic P-cycling experiments with fresh activated sludge from four full-scale EBPR plants. Since the specific carbon preferences for *Dechloromonas* were not known, a mixture of acetate, glucose, and casamino acids was used as carbon source during the anaerobic phase to accommodate a large range of potential requirements. Pure culture and *in situ* studies of *Dechloromonas* species has shown they can grow on volatile fatty acids and casamino acids [17], while well-known PAOs, such as *Ca*. Accumulibacter and *Tetrasphaera*, use acetate, amino acids, or sugars as substrates for P release under anaerobic conditions [2, 49, 50]. The addition of substrate stimulated an anaerobic release of ortho-P during the 3 h anaerobic phase (SFigure 1), as is typically seen for activated sludge from EBPR plants.

*In situ* identification and quantification of intracellular storage products using FISH-Raman microspectroscopy was performed on *Ca*. D. phosphatis in all four WWTPs. For *Ca*. D. phosphovora, lower abundances in the other three WWTPs limited FISH-Raman quantification to Viby WWTP. In all cases both species contained all three storage polymers (poly-P, glycogen, and PHA), and they both exhibited dynamic cycling of the polymers. In accordance with the accepted metabolic model for conventional PAOs as exemplified by *Ca*. Accumulibacter [49], the amount of intracellular poly-P was lowest after the anaerobic phase and highest after the aerobic phase, with some variations between plants (Figure 4A, Table 2). The highest values measured *in situ* for *Ca*. D. phosphatis was around 6.5*10^−14^g P cell^−1^ while *Ca*. D. phosphovora contained 6.3*10^−14^ g P cell^−1^ (Table 2). These values are slightly lower than for *Ca*. Accumulibacter (5-10*10^−14^ g P cell^−1^) but higher than for *Tetrasphaera* (1-2*10^−14^ g P cell^−1^). Their cell size seemed to correlate with poly-P content, with average *Dechloromonas* biovolume (2.35 µm^3^) lower than *Ca*. Accumulibacter (3.14 µm^3^), but higher than *Tetrasphaera* (0.45 µm^3^) [51].

**Table 2.** Summary of all storage compounds investigated in FISH-defined *Dechloromonas* cells in SBR reactor and EBPR plants.

| Sample type | Aerobic phase |  |  | Anaerobic phase |  |  |
| --- | --- | --- | --- | --- | --- | --- |
| | Poly-P<br>($\times 10^{-14}$ gP cell$^{-1}$) | PHA<br>($\times 10^{-14}$ gC cell$^{-1}$) | Glycogen<br>($\times 10^{-14}$ gC cell$^{-1}$) | Poly-P<br>($\times 10^{-14}$ gP cell$^{-1}$) | PHA<br>($\times 10^{-14}$ gC cell$^{-1}$) | Glycogen<br>($\times 10^{-14}$ gC cell$^{-1}$) |
| ***Candidatus Dechloromonas*** |  |  |  |  |  |  |
| **phosphatis** |  |  |  |  |  |  |
| EBPR plant Aalborg West* | 6.40 ± 0.23 | 3.14 ± 0.28 | 6.99 ± 0.17 | 0.71 ± 0.36 | 20.2 ± 0.14 | 1.21 ± 0.16 |
| EBPR plant Ejby-Mølle* | 6.95 ± 0.44 | 3.79 ± 0.24 | 7.37 ± 0.13 | 0.86 ± 0.33 | 23.2 ± 0.36 | 1.37 ± 0.34 |
| EBPR plant Lynetten* | 6.52 ± 0.18 | 3.22 ± 0.17 | 7.61 ± 0.25 | 0.65 ± 0.25 | 22.3 ± 0.13 | 1.64 ± 0.16 |
| EBPR plant Viby* | 6.21 ± 0.22 | 3.41 ± 0.23 | 7.48 ± 0.28 | 0.73 ± 0.36 | 21.3 ± 0.23 | 1.55 ± 0.39 |
| SBR reactor/casein hydrolysate | 6.75 ± 0.25 | 3.74 ± 0.36 | 7.25 ± 0.13 | 0.54 ± 0.14 | 21.8 ± 0.17 | 1.27 ± 0.27 |
| SBR reactor/acetate | | | | 0.88 ± 0.33 | 20.2 ± 0.41 | 1.31 ± 0.13 |
| SBR reactor/glucose | | | | 1.37 ± 0.25 | 19.2 ± 0.47 | 1.53 ± 0.37 |
| SBR reactor/glycine | | | | 2.65 ± 0.23 | 17.0 ± 0.43 | 1.94 ± 0.39 |
| SBR reactor/glutamate | | | | 2.54 ± 0.26 | 16.1 ± 0.36 | 1.81 ± 0.15 |
| SBR reactor/no substrates | | | | 5.45 ± 0.58 | 7.03 ± 0.32 | 6.05 ± 0.18 |
| ***Candidatus Dechloromonas*** |  |  |  |  |  |  |
| **phosphovora** |  |  |  |  |  |  |
| EBPR plant Viby* | 6.30 ± 0.27 | 3.52 ± 0.15 | 7.01 ± 0.38 | 0.99 ± 0.38 | 21.2 ± 0.28 | 1.33 ± 0.36 |
| SBR reactor/casein hydrolysate | 6.61 ± 0.23 | 3.63 ± 0.24 | 7.16 ± 0.24 | 0.61 ± 0.14 | 20.1 ± 0.42 | 1.07 ± 0.22 |
| SBR reactor/acetate | | | | 0.97 ± 0.36 | 21.1 ± 0.15 | 1.23 ± 0.17 |
| SBR reactor/glucose | | | | 1.46 ± 0.29 | 19.3 ± 0.16 | 1.45 ± 0.34 |
| SBR reactor/glycine | | | | 2.22 ± 0.13 | 17.9 ± 0.25 | 1.87 ± 0.17 |
| SBR reactor/glutamate | | | | 2.15 ± 0.22 | 18.0 ± 0.11 | 1.81 ± 0.24 |
| SBR reactor/no substrates | | | | 5.25 ± 0.56 | 7.38 ± 0.33 | 5.92 ± 0.13 |

\* EBPR biomass was fed with a mixture of acetate, glucose, and casamino acids.

**Figure 4.**
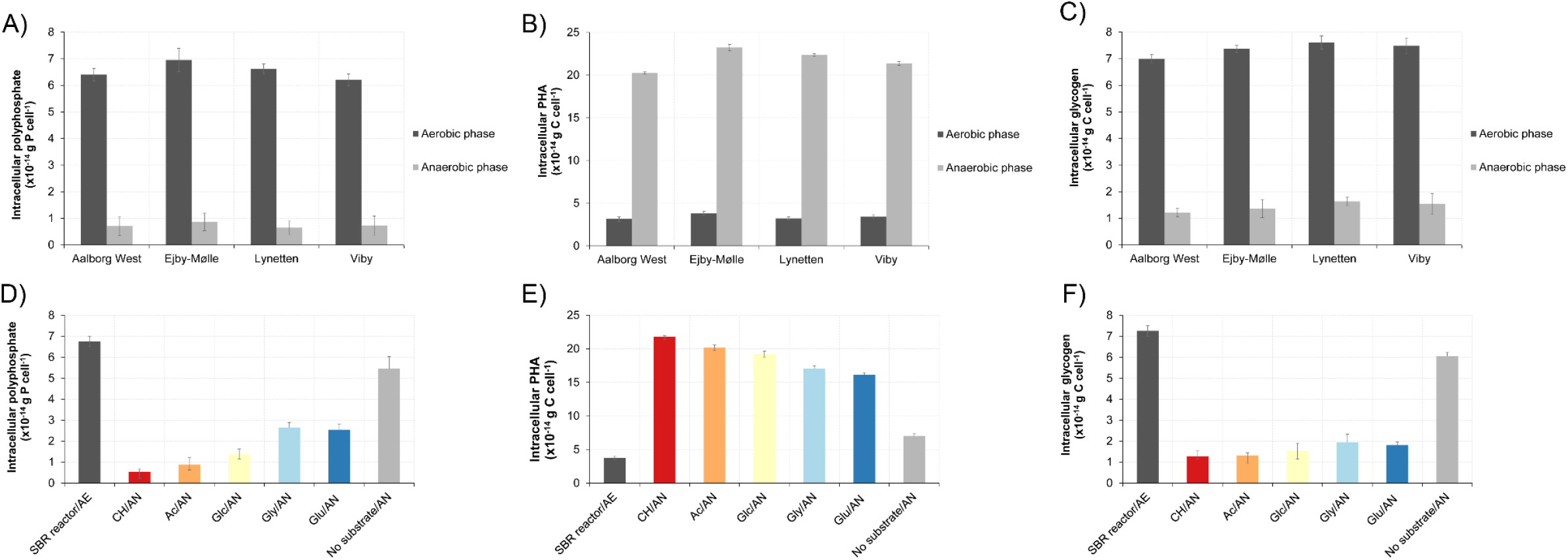
Dynamics of storage polymers measured by Raman microspectroscopy in FISH-defined cells of *Ca*. D. phosphatis by end of aerobic and anaerobic phases. A) Poly-P levels in *Ca*. D. phosphatis in different full-scale EBPR plants. B) PHA dynamics in *Ca*. D. phosphatis in different full-scale EBPR plants. C) Glycogen dynamics in *Ca*. D. phosphatis in different full-scale EBPR plants. D) Poly-P dynamics in *Ca*. D. phosphatis in SBR reactor fed with different substrates. E) PHA dynamics in *Ca*. D. phosphatis in SBR reactor fed with different substrates. F) Glycogen dynamics in *Ca*. D. phosphatis in SBR reactor fed with different substrates. SBR reactor is the initial sample collected at the end of the aerobic phase. CH = casein hydrolysate, Ac = acetate, Glc = glucose, Gly = glycine, Glu = glutamate. The sample indicated as “no substrate” has not been amended with additional substrate during anaerobic phase (control). AE = aerobic. AN = anaerobic.

The PHA content was also dynamic, with a build up during the anaerobic phase, reaching a similar level in all plants of approx. 21.2*10^−14^ g C cell^−1^ (Figure 4B, Table 2). Similarly, the glycogen level was reduced from approx. 7.2*10^−14^ in the aerobic state to approx. 1.3*10^−14^ g C cell^−1^ in the end of anaerobic phase (Figure 4C, Table 2). These conversions likely reflect the hydrolysis of glycogen used as an energy source for PHA formation under anaerobic conditions and its replenishment in the aerobic phase. The levels of PHA and glycogen were slightly lower than values recorded for *Ca*. Accumulibacter (30-39*10^−14^ g C cell^−1^ and 31-43*10^−14^ g C cell^−1^ for PHA and glycogen, respectively) most likely due to its smaller size [51]. In both *Dechloromonas* species, the C/P and C/C molar ratio were within the range of 0.3-0.4 that are reported for PAOs in previous studies [52, 53]. Our findings show that both *Dechloromonas* species expressed a phenotype very similar to the canonical PAO metabolism as described by *Ca*. Accumulibacter, with dynamic levels of poly-P, PHA, and glycogen under feast-famine conditions.

### Dynamics of storage compounds in *Dechloromonas* spp. fed with different substrates

To further investigate the physiology and the *in situ* dynamics of intracellular storage compounds in probe-defined *Ca*. D. phosphatis and *Ca*. D. phosphovora cells, feast-famine cycling experiments were carried out with biomass from a lab-scale SBR reactor. The reactor was conducted to enrich for novel PAOs (anaerobic/aerobic cycles with casein hydrolysate organic substrate added during the anaerobic phase) and contained 1-2% *Dechloromonas* spp. as quantified by qFISH. Different carbon sources (casein hydrolysate, acetate, glutamate, glycine, or glucose) were tested to investigate the potential to induce anaerobic poly-P hydrolysis and P release, indicating uptake of that particular substrate and conversion to PHA (Table 2). Addition of various substrates under anaerobic conditions was followed by a release of P and a subsequent uptake of P during aerobic conditions (SFigures 2-3). Both *Dechloromonas* species had in the end of the aerobic phase intracellular poly-P content in the range of 6.7-7.0*10^−14^ g P cell^−1^ (Figure 4D, Table 2), which is the same range as observed in the four full-scale EBPR plants tested. All substrates induced anaerobic degradation of intracellular poly-P with casein hydrolysate and acetate giving the highest release for both *Dechloromonas* species, and the two specific amino acids glycine and glutamate the lowest (Figure 4D, Table 2 and SFigure2). Glucose induced P-release from the biomass and also intracellular dynamics of poly-P, PHA, and glycogen, indicating uptake and conversion of glucose by *Dechloromonas* species. However, glucose uptake is not observed in *Dechloromonas* pure cultures [17] and is also not encoded by MAGs representing the two species (see below). Thus, the observed glucose pattern may be due to uptake and metabolism by other members of the community, which then provide substrates that can be used by the *Dechloromonas* species. Intracellular cycling of PHA and glycogen occurred in accordance with the results from the full-scale biomass investigation (Figure 4E-F, Table 2), and with similar C/P and C/C ratios of 0.3-0.4.

Interestingly, these substrate uptake patterns of *Dechloromonas* species, also observed by Qiu et al., [50], indicate a niche partially overlapping with the two other common PAOs, *Ca*. Accumulibacter with acetate as primary substrate [49, 54, 55], and *Tetrasphaera* with amino acids and glucose as primary substrates [2, 56], although some *Ca*. Accumulibacter may also use amino acids [49, 54, 55]. Since their ecophysiology is very similar to *Ca*. Accumulibacter, former “PAO” studies without reliable identification of all PAOs may have misinterpreted their findings and wrongly assumed the P-dynamics to be due to *Ca*. Accumulibacter.

### Metabolic potential of uncultivated *Dechloromonas* spp. in EBPR systems

In order to look further into the potential physiology of the *Dechloromonas* genus in activated sludge, 10 *Dechloromonas* MAGs obtained by Singleton et al. [26] from Danish EBPR plants were annotated and the potential for particular pathways important for the PAO metabolism were identified (Figures 5-6). The MAGs represented 7 species, where *Ca*. D. phosphatis and *Ca*. D. phosphovora (and ASV85) were represented by the MAGs Ribe_DB9 and Skiv_DB10, respectively (Figures 4-5). None of these MAGs possessed >95% average nucleotide identity to the isolates, so they all represent novel uncultured species.

**Figure 5.**
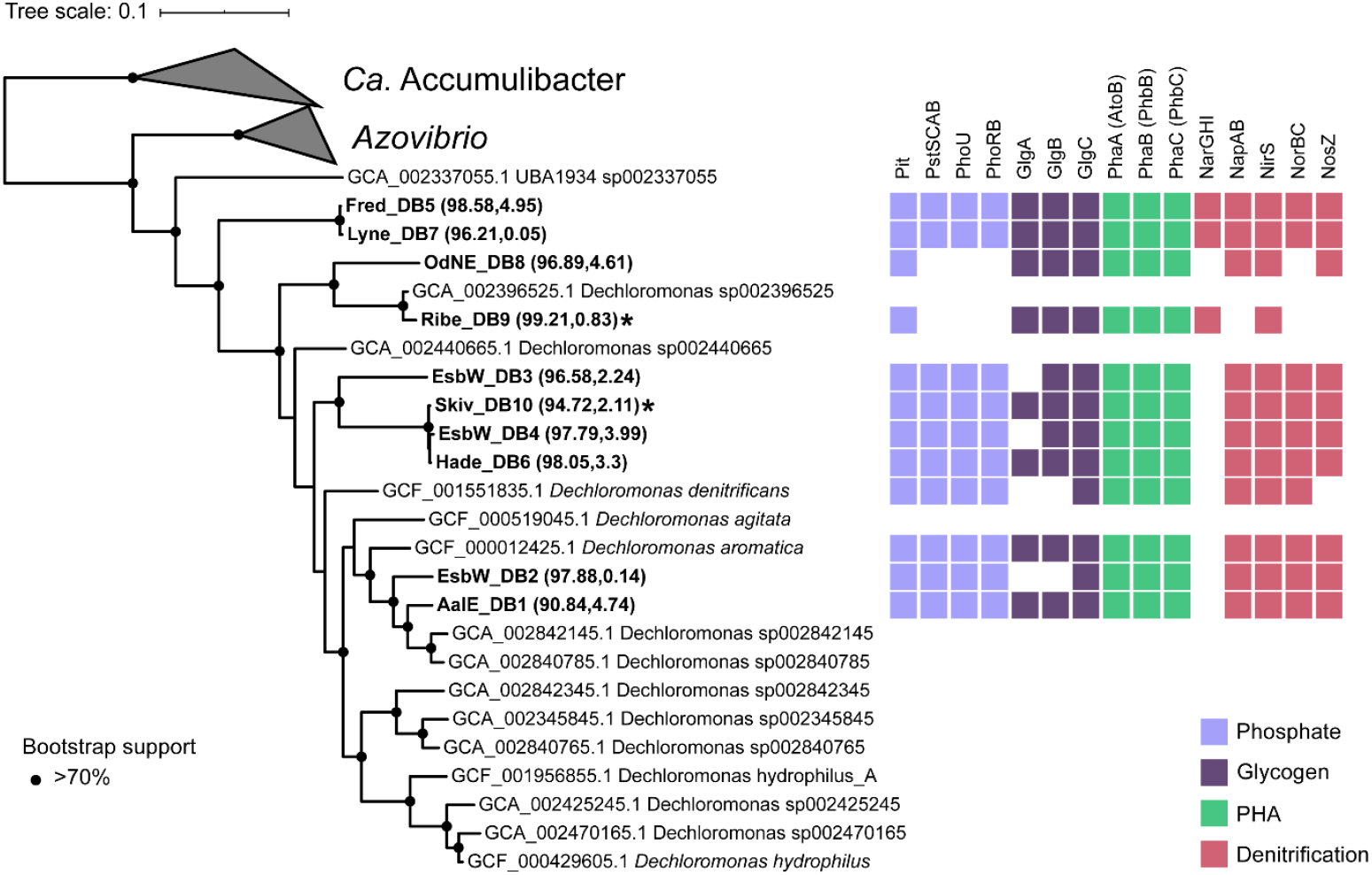
Maximum likelihood tree created using the WAG + GAMMA model in IQ-TREE and the alignment of 120 concatenated proteins created by GTDB-Tk with GTDB release89. Five *Ca*. Accumulibacter genomes were used as the outgroup. Genome completeness and contamination % estimates for the MAGs are indicated within the parentheses. Genomes in bold indicate the MAGs recovered from the Danish WWTPs. *Ca*. D. phosphatis and *Ca*. D. phosphovora are represented by the MAGs Ribe_DB9 and Skiv_DB10 respectively, indicated by asterisks. Bootstrap support >70% is indicated by the solid black circles. NosZ was not identified in the *D. denitrificans* genome, however, it is present in NCBI under accession number KT592356.1.

**Figure 6.**
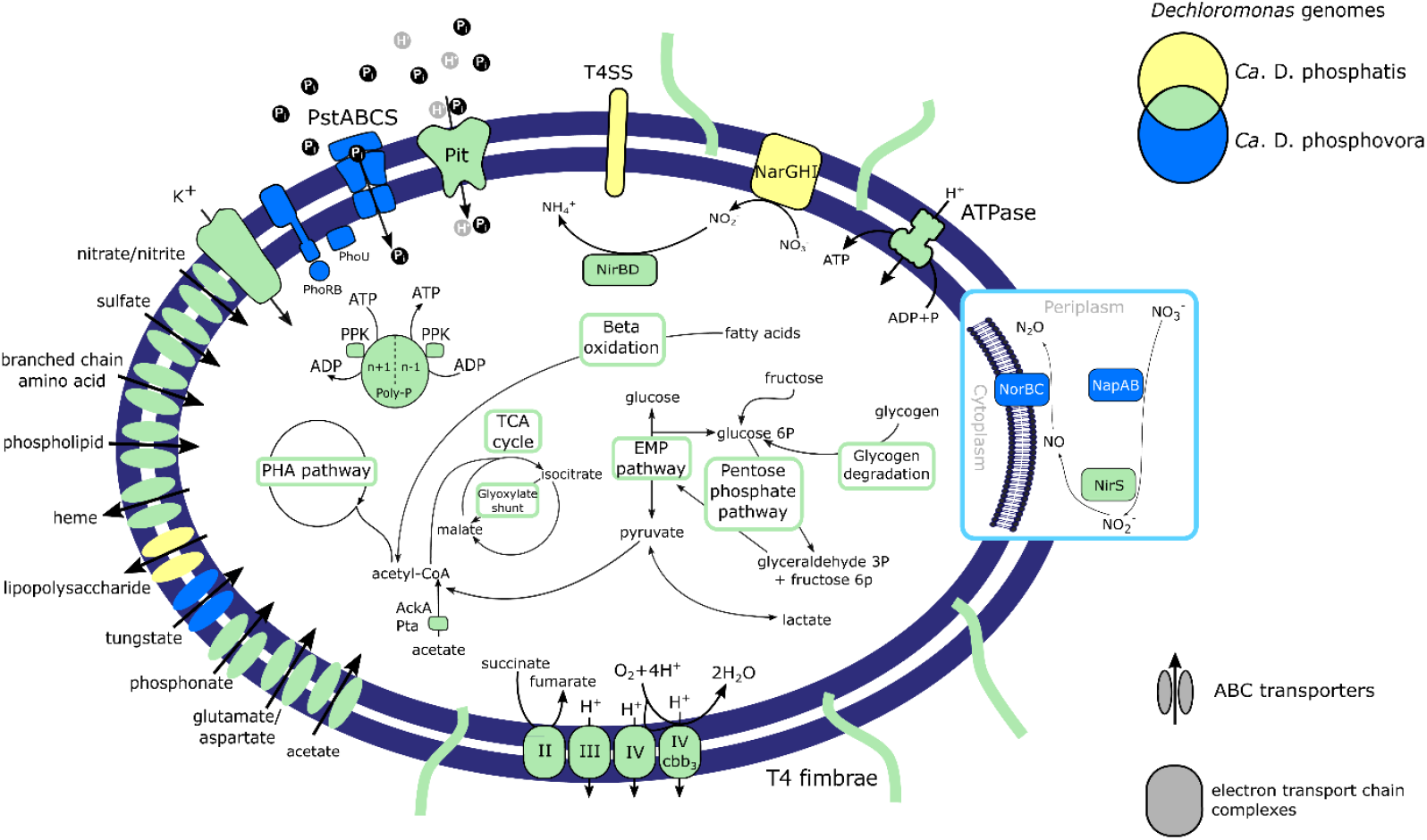
Metabolic model of the two most abundant *Dechloromonas* species. Colours indicate the species or combination of species (Venn diagram) that encode the potential for the enzyme or cycle. Abbreviation: TCA = tricarboxylic acid cycle, EMP = Embden-Meyerhof-Parnas pathway (glycolysis), PHA = polyhydroxyalkanoate pathway, I = complex I NADH dehydrogenase, II = complex II succinate dehydrogenate, III = complex III cytochrome bc1, IV = cytochrome c oxidase, IV cbb3 = complex IV cytochrome cbb3 oxidase, Pit = inorganic phosphate transporter family, PstSCAB = inorganic phosphate ABC transporter, PhoRB = two component system for phosphate regulation, PhoU = phosphate transport system accessory protein, Poly-P = polyphosphate, Ppk = polyphosphate kinase, T4SS = type IV secretion system, NapAB = periplasmatic nitrate reductase, NarGHI = membrane-bound nitrate reductase, NirS = nitrite reductase, NorBC = nitric oxide reductase, AckA = acetate kinase, Pta = phosphotransacetylase.

Metabolic reconstruction of the *Dechloromonas* species MAGs confirmed the presence of central carbon pathways, including the glycolysis, TCA cycle, and glyoxylate pathways, along with genes essential for PHA and glycogen accumulation (Figures 5-6 and STable1). We confirmed the genomic potential for uptake of two carbon sources used in the P-release experiments by identifying the transporters for acetate and glutamate (Figure 6). However, glycine (*cyc*A) and glucose transporter genes could not be detected in the genomes, suggesting that other members of the community processed these substrates into components that could be used by the *Dechloromonas* species in the SBR experiment. The identification of both glycine hydroxymethyltransferase (*gly*A) and glutamate dehydrogenase (*gdh*A) in the MAGs suggests that the *Dechloromonas* species processed these amino acids to pyruvate and 2-oxoglutarate, respectively, for integration into central carbon metabolism. Interestingly, all MAGs encoded the complete pathway for PHA accumulation, also confirmed in the genomes of the two closely related pure cultures, *Dechloromonas agitata* and *Dechloromonas denitrificans. Ca*. D. phosphatis, and *Ca*. D. phosphovora encoded the genes for glycogen accumulation, supporting the experimental results. Two of the other MAGs, representing different *Dechloromonas* spp. (EsbW_DB3 and EsbW_DB2), lacked the genes for glycogen synthase (*glgA*) and 1,4-alpha-glucan branching enzyme (*glgB*) for the glycogen accumulation pathway (Figures 5-6 and STable1). These genes were also absent in the isolate *D. denitrificans*. Other carbon metabolisms encoded by the MAGs included the potential for beta-oxidation of fatty acids, and fructose degradation (STable1 and SDataFile1). Lactate (both isomers for Ribe_DB9, D-lactate only for others) may be used as an additional electron donor and carbon source, as in *D. denitrificans* [17], or could be produced as a product of fermentation of pyruvate to lactate, depending on the prevailing environmental conditions.

Although the potential for polyphosphate metabolism was present in all the MAGs and in *D. denitrificans* and *D. aromatica*, a key difference was found in the phosphate transporters. While the majority of the MAGs possessed the high-affinity PstSCAB transporter system, the most abundant *Ca*. D. phosphatis (Ribe_DB9) and its close relative OdNE_DB8 possessed only the low-affinity Pit transporters (Figure 5). This supports the proposal that Pit is the phosphate transporter vital to the PAO phenotype [57]. The Pit system is essential as it generates a proton motive force (PMF) by the export of metal-phosphates in symport with a proton under anaerobic conditions, which seems to drive volatile fatty acids (VFA) uptake in *Ca*. Accumulibacter [57, 58]. Moreover, the high-affinity PstSCAB transporter system may not be essential in a P-rich environment such as activated sludge. Further analysis, for example with the aid of transcriptomics, may be useful to determine the relevance and usage of these transporters in different conditions. A potential GAO phenotype *in situ* was previously suggested for probe-defined *Dechloromonas* (likely *Ca*. Dechloromonas phosphovora) [22], but GAOs typically encode only the high affinity PstSCAB uptake system [57]. The presence of genes encoding the Pit system, supported by the experimental poly-P accumulation evidence, is in accordance with a conventional PAO metabolism. It is possible that *Dechloromonas* species, as previously shown for *Ca*. Accumulibacter [59, 60], can exhibit different metabolisms, spanning from polyphosphate-to glycogen-based phenotypes depending on the environmental conditions, but additional analysis is needed to verify this potential metabolic versatility.

Nitrogen metabolism also varied between the different MAGs. All the MAGs presented the potential for dissimilatory nitrate reduction and some of them (EsbW_DB2, Fred_DB5, and Lyne_DB7) possessed the genes for nitrogen fixation (Figures 5-6 and STable 1). The majority of MAGs (8/10) encoded the potential for complete denitrification, while *Ca*. D. phosphatis (Ribe_DB9) and OdNE_DB8 only encoded reduction of nitrate to nitric oxide (Figures 5-6). For OdNE_DB8, the absence of the nitric oxide reductase (NorBC) may be due to incompleteness, as this genome contained all other components of the denitrification pathway, including the nitrous oxide reductase (NosZ). The *Ca*. D. phosphatis (Ribe_DB9) genome did not encode the periplasmic nitrate reductase NapAB, present in all other recovered genomes, but the respiratory nitrate reductase NarGHI (which is also present in combination with NapAB in Fred_DB5 and Lyne_DB7). This could indicate a difference in their ecological niche, as the Nap enzyme is usually not involved in anaerobic respiration [61], while the presence of a respiratory nitrate reductase may allow *Ca*. D. phosphatis (Ribe_DB9) to dominate under anoxic conditions. Previous studies on *Ca*. Accumulibacter species [61–63] showed that the Nar enzyme is essential for anoxic phosphorus uptake using nitrate, and its absence in the other MAGs may indicate an inability to use this metabolic function. The subsequent steps of the denitrification pathway also varied between the different MAGs. All of them encoded a dissimilatory nitrite reductase (NirS), while nitric oxide reductase (NosZ) was found in all the MAGs, excluding *Ca*. D. phosphatis (Ribe_DB9). As this MAG lacked other genes of the denitrification pathway (Figures 5-6, STable1), important for the reduction of toxic intermediates [64], this organism most likely lost the metabolic capability to use nitric oxide and nitrous oxide as terminal electron acceptors. Additional *in situ* studies are needed to verify this metabolic trait and its possible effects on niche partitioning within the genus and with other important PAOs.

### Taxonomic proposal

As no axenic cultures representing the novel species are available, we propose the names *Candidatus* Dechloromonas phosphatis sp. nov. [Phos.pha’tis. phosphatis from L. nm. *phosphas*, phosphate: a microorganism which can accumulate polyphosphate] and *Candidatus* Dechloromonas phosphovora [Phos.pho’vo.ra, from L. nm. *phosphorus*, phosphorus; L. v. *vora*, devour: phosphorus-accumulating microorganism] for the two species, based on the recommendations by Murray and Stackebrandt [65]. The formal proposal of the new species is given in STables3-4.

### Concluding remarks and future perspectives

Here, we have provided the first experimental and genomic insights into the physiology of uncultivated *Dechloromonas* spp. We have shown that these lineages are PAOs that are often abundant, and actively involved in P removal in full-scale EBPR plants, and they should be regarded as important PAOs along with *Ca*. Accumulibacter and *Tetrasphaera*. They exhibit a physiology very similar to *Ca*. Accumulibacter, but seem more diverse in their substrate utilization profile and share some traits with *Tetrasphaera* too. They may also be important for nitrogen removal with most species possessing full denitrifying capabilities. The co-occurrence of these three genera in most EBPR plants suggests niche differentiation and functional redundancy, which may be even more pronounced by the presence of other, so far undescribed PAOs. Some putative PAOs can occasionally be abundant in specific plants [15] and our recent discovery of a new putative PAO, *Ca*. Methylophosphatis [26], strongly suggests there are still several undescribed PAOs. This has important implications for the study of EBPR communities, as they are critical to nutrient cycling in many wastewater treatment plants and warrant specific attention as societies transition to improve resource recovery from these systems.

## Supporting information

SDataFile1

SDataFile2

STable1

## Acknowledgements

The project was funded by the Villum Foundation (Dark Matter grant13351) and Aalborg University.

## SUPPLEMENTARY MATERIAL

**SFigure 1.**
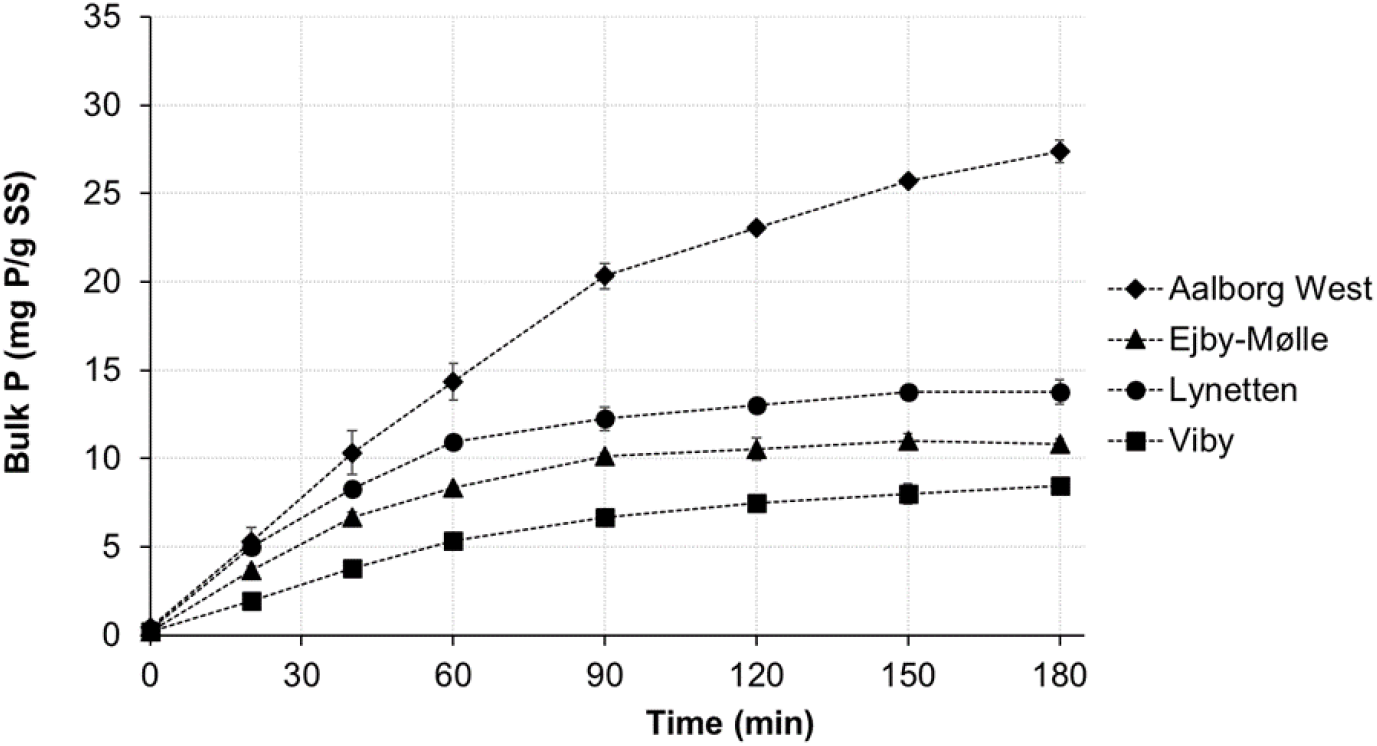
Anaerobic phosphate release profiles from 4 full-scale EBPR plant activated sludge samples, amended with a mixture of acetate, glucose, and casamino acids with a final concentration of 500, 250, and 250 mg L^-1^, respectively.

**SFigure 2.**
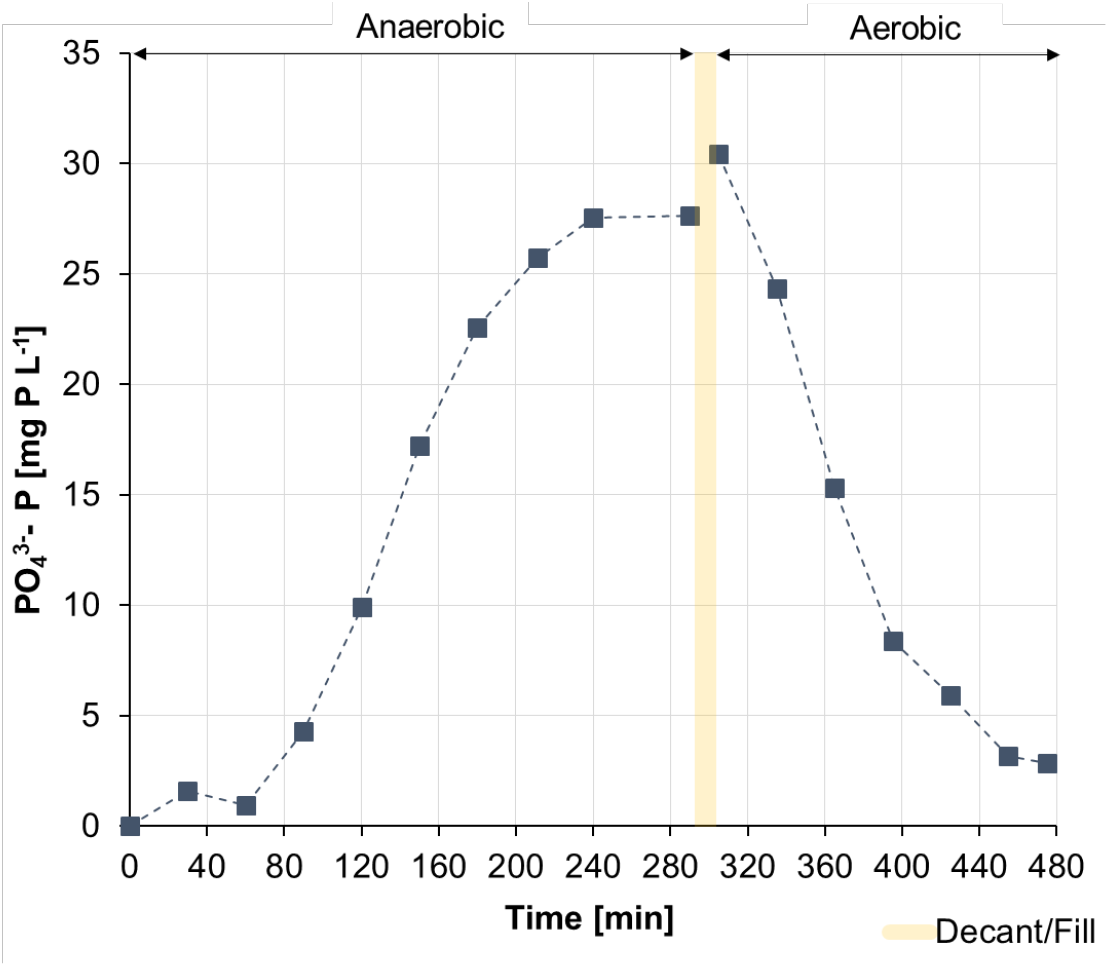
Phosphate release and phosphate uptake from SBR biomass containing PAOs during an operational cycle. The substrate added under anaerobic conditions was casein hydrolysate at an initial concentration of 200 mg COD L^-1^

**SFigure 3.**
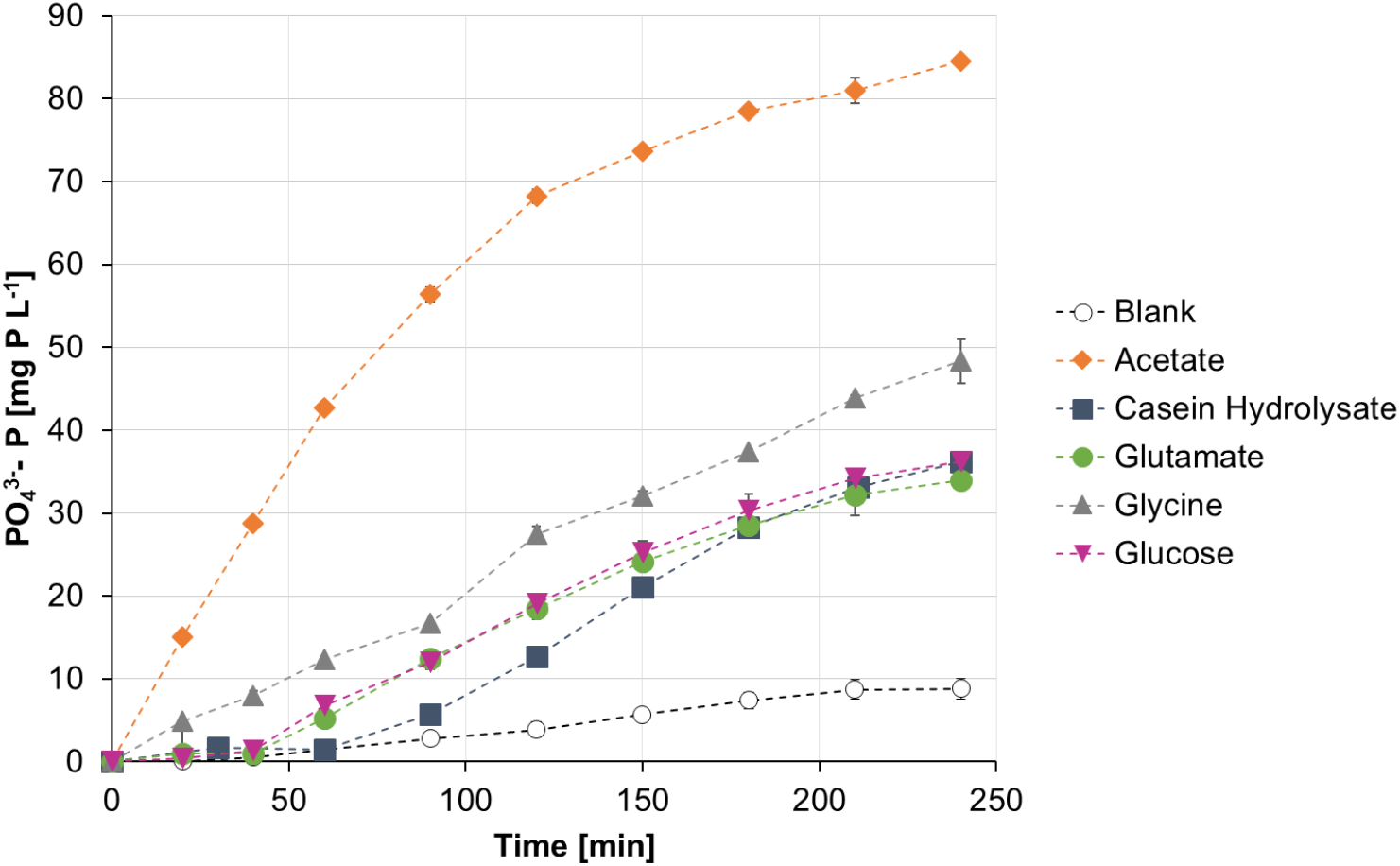
Anaerobic phosphate release profiles obtained with different carbon sources with an initial cycle concentration of 200 mg COD L^-1^ from SBR biomass containing PAOs, including *Dechloromonas*.

SDataFile1: description (100% complete KO modules identified using EnrichM)

SDataFile2: description (16S rRNA genes)

**STable 2.**
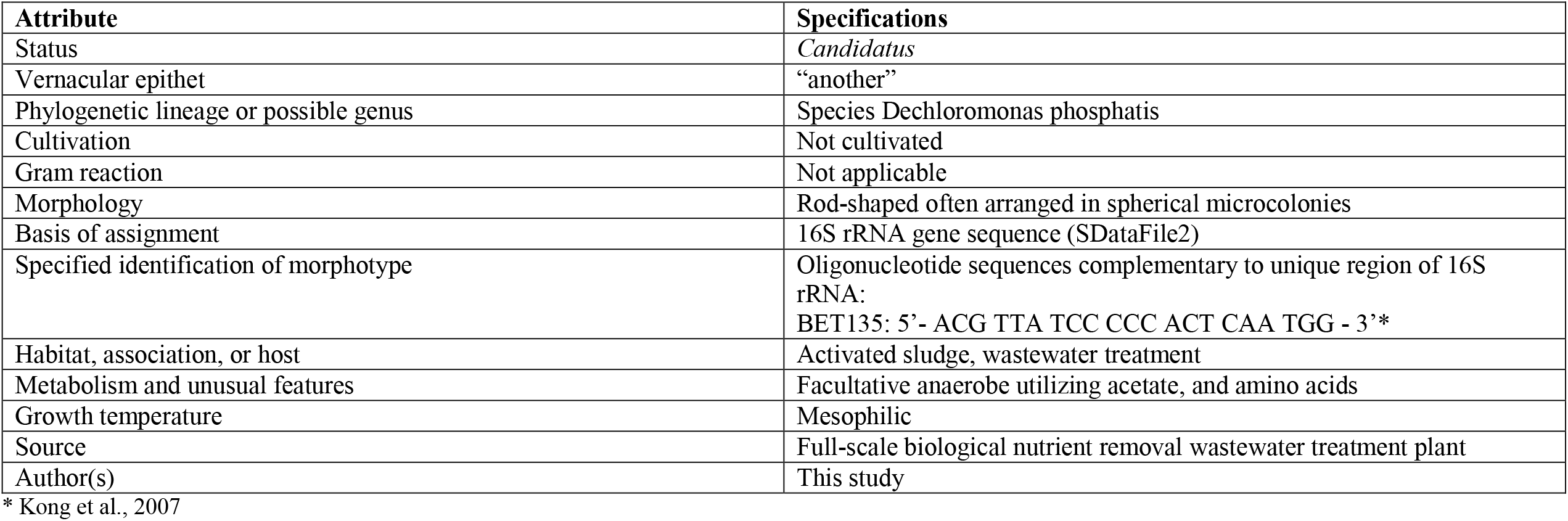
Items for inclusion in the codified record of provisional taxon for *Candidatus* Dechloromonas phosphatis

**STable 3.**
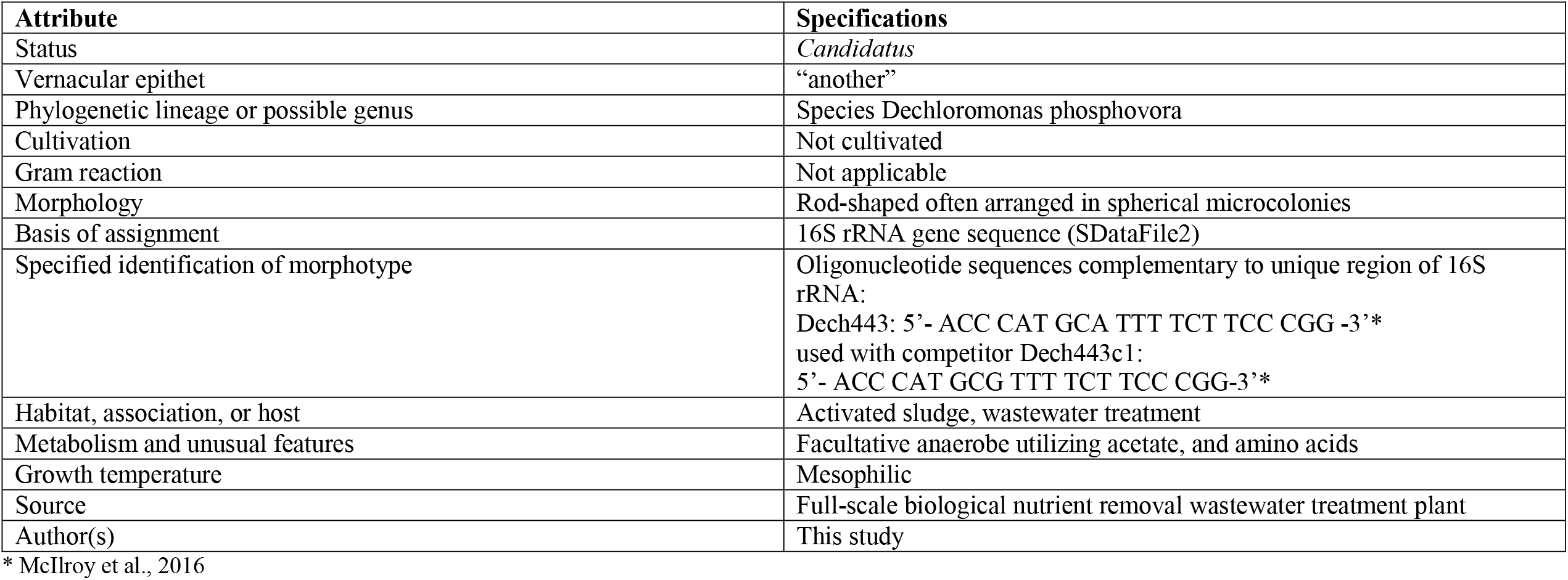
Items for inclusion in the codified record of provisional taxon for *Candidatus* Dechloromonas phosphovora

## References

1. Nielsen PH, Mcilroy SJ, Albertsen M, Nierychlo M. Re-evaluating the microbiology of the enhanced biological phosphorus removal process. Curr Opin Biotechnol 2019; 57: 111–118.

2. Marques R, Santos J, Nguyen H, Carvalho G, Noronha JP, Nielsen PH, et al. Metabolism and ecological niche of Tetrasphaera and Ca. Accumulibacter in enhanced biological phosphorus removal. Water Res 2017; 122: 159–171.

3. Camejo PY, Oyserman BO, McMahon KD, Noguera DR. Integrated Omic Analyses Provide Evidence that a “Candidatus Accumulibacter phosphatis” Strain Performs Denitrification under Microaerobic Conditions. mSystems 2019; 4: 1–23.

4. Oyserman BO, Noguera DR, Del Rio TG, Tringe SG, McMahon KD. Metatranscriptomic insights on gene expression and regulatory controls in Candidatus Accumulibacter phosphatis. ISME J 2016; 10: 810–822.

5. Tu Y, Schuler AJ. Low acetate concentrations favor polyphosphate-accumulating organisms over glycogen-accumulating organisms in enhanced biological phosphorus removal from wastewater. Environ Sci Technol 2013; 47: 3816–3824.

6. Marques R, Ribera-guardia A, Santos J, Carvalho G, Reis MAM, Pijuan M, et al. Denitrifying capabilities of Tetrasphaera and their contribution towards nitrous oxide production in enhanced biological phosphorus removal processes. Water Res 2018; 137: 262–272.

7. Fernando EY, McIlroy SJ, Nierychlo M, Herbst F-A, Petriglieri F, Schmid MC, et al. Resolving the individual contribution of key microbial populations to enhanced biological phosphorus removal with Raman–FISH. ISME J 2019; 1933–1946.

8. Kawaharasaki M, Tanaka H, Kanagawa T, Nakamura K. In situ identification of polyphosphate-accumulating bacteria in activated sludge by dual staining with rRNA-targeted oligonucleotide probes and 4’,6-diaimidino-2-phenylindol (DAPI) at a polyphosphate-probing concentration. Water Res 1999; 33: 257–265.

9. Crocetti GR, Hugenholtz P, Bond PL, Schuler AJ, Keller J, Jenkins D, et al. Identification of polyphosphate-accumulating organisms and design of 16SrRNA-directed probes for their detection and quantitation. Appl Environ Microbiol 2000; 66: 1175–1182.

10. Kong Y, Nielsen JL, Nielsen PH. Identity and Ecophysiology of Uncultured Actinobacterial Polyphosphate-Accumulating Organisms in Full-Scale Enhanced Biological Phosphorus Removal Plants. Appl Environ Microbiol 2005; 71: 4076–4085.

11. Kong Y, Xia Y, Nielsen JL, Nielsen PH. Structure and function of the microbial community in a full-scale enhanced biological phosphorus removal plant. Microbiology 2007; 153: 4061–4073.

12. Goel R, Sanhueza P, Noguera D. Evidence of Dechloromonas sp. participating in enhanced biological phosphorous removal (EBPR) in a bench-scale aerated-anoxic reactor. Proc Water Environ Fed 2005; 41: 3864–3871.

13. Terashima M, Yama A, Sato M, Yumoto I, Kamagata Y, Kato S. Culture-Dependent and - independent Identification of Polyphosphate-Accumulating Dechloromonas spp. Predominating in a Full-Scale Oxidation Ditch Wastewater Treatment Plant. Microbes Environ Environ 2016; 31: 449–455.

14. Wang B, Jiao E, Guo Y, Zhang L, Meng Q, Zeng W, et al. Investigation of the polyphosphate-accumulating organism population in the full-scale simultaneous chemical phosphorus removal system. Environ Sci Pollut Res 2020; 27: 37877–37886.

15. Stokholm-Bjerregaard M, McIlroy SJ, Nierychlo M, Karst SM, Albertsen M, Nielsen PH. A critical assessment of the microorganisms proposed to be important to enhanced biological phosphorus removal in full-scale wastewater treatment systems. Front Microbiol 2017; 8: 718.

16. Achenbach LA, Michaelidou U, Bruce RA, Fryman J, Coates JD. Dechloromonas agitata gen. nov., sp. nov. and Dechlorosoma suillum gen. nov., sp. nov., two novel environmentally dominant (per)chlorate-reducing bacteria and their phylogenetic position. Int J Syst Evol Microbiol 2001; 51: 527–533.

17. Horn MA, Ihssen J, Matthies C, Schramm A, Acker G, Drake HL, et al. Dechloromonas denitrificans sp. nov., Flavobacterium denitrificans sp. nov., Paenibacillus anaericanus sp. nov. and Paenibacillus terrae strain MH72, N2O-producing bacteria isolated from the gut of the earthworm Aporrectodea caliginosa. Int J Syst Evol Microbiol 2005; 55: 1255–1265.

18. Acevedo B, Murgui M, Borrás L, Barat R. New insights in the metabolic behaviour of PAO under negligible poly-P reserves. Chem Eng J 2017; 311: 82–90.

19. Yuan Y, Liu J, Ma B, Liu Y, Wang B, Peng Y. Improving municipal wastewater nitrogen and phosphorous removal by feeding sludge fermentation products to sequencing batch reactor (SBR). Bioresour Technol 2016; 222: 326–334.

20. Lv X, Shao M, Li C, Li J, Gao X, Sun F. A Comparative Study of the Bacterial Community in Denitrifying and Traditional Enhanced Biological Phosphorus Removal Processes. Microbes Environ 2014; 29: 261–268.

21. Salinero KK, Keller K, Feil WS, Feil H, Trong S, Bartolo G Di, et al. Metabolic analysis of the soil microbe Dechloromonas aromatica str. RCB anaerobic pathways for aromatic degradation. BMC Genomics 2009; 23: 1–23.

22. McIlroy SJ, Starnawska A, Starnawski P, Saunders AM, Nierychlo M, Nielsen PH, et al. Identification of active denitrifiers in full-scale nutrient removal wastewater treatment systems. Environ Microbiol 2016; 18: 50–64.

23. Hesselsoe M, Fu S, Schloter M, Bodrossy L, Iversen N, Roslev P, et al. Isotope array analysis of Rhodocyclales uncovers functional redundancy and versatility in an activated sludge. ISME J 2009; 3: 1349–1364.

24. Ahn J, Schroeder S, Beer M, McIlroy S, Bayly RC, May JW, et al. Ecology of the Microbial Community Removing Phosphate from Wastewater under Continuously Aerobic Conditions in a Sequencing Batch Reactor. Appl Environ Microbiol 2007; 73: 2257–2270.

25. Dueholm MS, Andersen KS, McIlroy SJ, Kristensen JM, Yashiro E, Karst SM, et al. Generation of comprehensive ecosystems-specific reference databases with species-level resolution by high-throughput full-length 16S rRNA gene sequencing and automated taxonomy assignment (AutoTax). bioRxiv 2020; 672873.

26. Singleton CM, Petriglieri F, Kristensen JM, Kirkegaard RH, Michaelsen TY, Andersen MH, et al. Connecting structure to function with the recovery of over 1000 high-quality activated sludge metagenome-assembled genomes encoding full-length rRNA genes using long-read sequencing. bioRxiv 2020; 2020.05.12.088096.

27. Albertsen M, Hugenholtz P, Skarshewski A, Nielsen KL, Tyson GW, Nielsen PH. Genome sequences of rare, uncultured bacteria obtained by differential coverage binning of multiple metagenomes. Nat Biotechnol 2013; 31: 533–8.

28. Parks DH, Rinke C, Chuvochina M, Chaumeil P, Woodcroft BJ, Evans PN, et al. Recovery of nearly 8,000 metagenome-assembled genomes substantially expands the tree of life. Nat Microbiol 2017; 2: 1533–1542.

29. Smolders GJF, Meij J Van Der, Loosdrecht MCM Van, Heijnen JJ. Model of the Anaerobic Metabolism of the Biological Phosphorus Removal Process: Stoichiometry and pH Influence. Biotechnol Bioeng 1994; 43: 461–470.

30. Jørgensen MK, Nierychlo M, Nielsen AH, Larsen P, Christensen ML, Nielsen PH. Unified understanding of physico-chemical properties of activated sludge and fouling propensity. Water Res 2017; 120: 117–132.

31. Nielsen JL. Protocol for fluorescence in situ hybridization (FISH) with rRNA-targeted oligonucleotides. FISH Handbook for Biological Wastewater Treatment. 2009. pp 73–84.

32. McIlroy SJ, Kirkegaard RH, McIlroy B, Nierychlo M, Kristensen JM, Karst SM, et al. MiDAS 2.0: An ecosystem-specific taxonomy and online database for the organisms of wastewater treatment systems expanded for anaerobic digester groups. Database 2017; 2017: 1–9.

33. Nierychlo M, Andersen KS, Xu Y, Green N, Jiang C, Albertsen M, et al. MiDAS 3: An ecosystem-specific reference database, taxonomy and knowledge platform for activated sludge and anaerobic digesters reveals species-level microbiome composition of activated sludge. Water Res 2020; 115955.

34. RStudio Team. RStudio: Integrated Development Environment for R. 2015. Boston, MA.

35. Albertsen M, Karst SM, Ziegler AS, Kirkegaard RH, Nielsen PH. Back to basics - The influence of DNA extraction and primer choice on phylogenetic analysis of activated sludge communities. PLoS One 2015; 10: 1–15.

36. Wickham H. ggplot2 - Elegant Graphics for Data Analysis. Springer. 2009. Springer Science & Business Media.

37. Ludwig W, Strunk O, Westram R, Richter L, Meier H, Yadhukumar a., et al. ARB: A software environment for sequence data. Nucleic Acids Res 2004; 32: 1363–1371.

38. Quast C, Pruesse E, Yilmaz P, Gerken J, Schweer T, Yarza P, et al. The SILVA ribosomal RNA gene database project: Improved data processing and web-based tools. Nucleic Acids Res 2013; 41: 590–596.

39. Yilmaz LS, Parnerkar S, Noguera DR. MathFISH, a web tool that uses thermodynamics-based mathematical models for in silico evaluation of oligonucleotide probes for fluorescence in situ hybridization. Appl Environ Microbiol 2011; 77: 1118–1122.

40. Daims H, Stoecker K, Wagner M. Fluorescence in situ hybridization for the detection of prokaryotes. In: Osborn AM, Smith CJ (eds). Molecular Microbial Ecology. 2005. Taylor & Francis, New York, pp 213–239.

41. Daims H, Lücker S, Wagner M. Daime, a Novel Image Analysis Program for Microbial Ecology and Biofilm Research. Environ Microbiol 2006; 8: 200–213.

42. Chaumeil P, Mussig AJ, Parks DH, Hugenholtz P. Genome analysis GTDB-Tk?: a toolkit toclassify genomes with the Genome Taxonomy Database. Bioinformatics 2019; 36: 1925–1927.

43. Olm MR, Brown CT, Brooks B, Banfield JF. dRep?: a tool for fast and accurate genomic comparisons that enables improved genome recovery from metagenomes through de-replication. ISME J 2017; 11: 2864–2868.

44. Kanehisa M, Sato Y, Kawashima M, Furumichi M, Tanabe M. KEGG as a reference resource for gene and protein annotation. Nucleic Acids Res 2016; 44: 457–462.

45. Seemann T. Prokka?: rapid prokaryotic genome annotation. Bioinformatics 2014; 30: 2068–2069.

46. Nawrocki EP, Kolbe DL, Eddy SR. Infernal 1.0: inference of RNA alignments. 2009; 25: 1335–1337.

47. Nguyen L, Schmidt HA, Haeseler A Von, Minh BQ. IQ-TREE: A Fast and Effective Stochastic Algorithm for Estimating Maximum-Likelihood Phylogenies. 2014; 32: 268–274.

48. Wolterink A, Kim S, Muusse M, Kim IS, Roholl PJM, Ginkel CG Van, et al. Dechloromonas hortensis sp. nov. and strain ASK-1, two novel (per)chlorate-reducing bacteria, and taxonomic description of strain GR-1. 2005; 1: 2063–2068.

49. Oehmen A, Lemos PC, Carvalho G, Yuan Z, Blackall LL, Reis MAM. Advances in enhanced biological phosphorus removal: From micro to macro scale. Water Res 2007; 41: 2271–2300.

50. Qiu G, Zuniga-montanez R, Law Y, Swa S. Polyphosphate-accumulating organisms in full-scale tropical wastewater treatment plants use diverse carbon sources. Water Res 2019; 149: 469–510.

51. Petriglieri F, Petersen JF, Peces M, Nierychlo M, Hansen K, Baastrand CE, et al. Quantification of biologically and chemically bound phosphorus in activated sludge from EBPR plants. Submitted.

52. Hesselmann RPX, Von Rummel R, Resnick SM, Hany R, Zehnder AJB. Anaerobic metabolism of bacteria performing enhanced biological phosphate removal. Water Res 2000; 34: 3487–3494.

53. Acevedo B, Oehmen A, Carvalho G, Seco A, Borrás L, Barat R. Metabolic shift of polyphosphate-accumulating organisms with different levels of polyphosphate storage. Water Res 2012; 6: 1889–1900.

54. Flowers JJ, He S, Malfatti S, Glavina T, Tringe SG, Hugenholtz P, et al. Comparative genomics of two ‘Candidatus Accumulibacter’ clades performing biological phosphorus removal. ISME J 2013; 7: 2301–2314.

55. Qiu G, Liu X, Saw NMMT, Law Y, Zuniga-Montanez R, Thi SS, et al. Metabolic Traits of Candidatus Accumulibacter clade IIF Strain SCELSE-1 Using Amino Acids As Carbon Sources for Enhanced Biological Phosphorus Removal. Environ Sci Technol 2019; 54: 2448–2458.

56. Kristiansen R, Thi H, Nguyen T, Saunders AM, Nielsen JL, Wimmer R, et al. A metabolic model for members of the genus Tetrasphaera involved in enhanced biological phosphorus removal. ISME J 2013; 7: 543–554.

57. McIlroy SJ, Albertsen M, Andresen EK, Saunders AM. ‘Candidatus Competibacter’-lineage genomes retrieved from metagenomes reveal functional metabolic diversity. ISME J 2014; 8: 613–624.

58. Saunders AM, Mabbett AN, Mcewan AG, Blackall LL. Proton motive force generation from stored polymers for the uptake of acetate under anaerobic conditions. FEMS Microbiol Lett 2007; 274: 245–251.

59. Erdal UG, Erdal ZK, Daigger GT, Randall CW. Is it PAO-GAO competition or metabolic shift in EBPR system?? Evidence from an experimental study. Water Sci Technol 2008; 58: 1329–1334.

60. Zhou Y, Pijuan M, Zeng RJ, Lu H, Ã ZY. Could polyphosphate-accumulating organisms (PAOs) be glycogen-accumulating organisms (GAOs)? Water Res 2008; 42: 2361–2368.

61. Weissbrodt DG, Lopez-vazquez CM, Welles L. “Candidatus Accumulibacter delftensis”?: A clade IC novel polyphosphate-accumulating organism without denitrifying activity on nitrate. Water Res 2019; 161: 136–151.

62. Camejo PY, Owen BR, Martirano J, Ma J, Kapoor V, Santo J, et al. Candidatus Accumulibacter phosphatis clades enriched under cyclic anaerobic and microaerobic conditions simultaneously use different electron acceptors. Water Res 2016; 102: 125–137.

63. Skennerton CT, Barr JJ, Slater FR, Bond PL, Tyson GW. Expanding our view of genomic diversity in Candidatus Accumulibacter clades. Environ Microbiol 2015; 17: 1574–1585.

64. Hendriks J, Oubrie A, Castresana J, Urbani A, Gemeinhardt S, Saraste M. Nitric oxide reductases in bacteria. Biochim Biophys Acta - Bioenerg 2000; 1459: 266–273.

65. Murray RGE, Stackebrandt E. Taxonomic Note?: Implementation of the Provisional Status Candidatus for Incompletely Described Procaryotes. Int J Syst Evol Microbiolology 1995; 45: 186–187.

